# pSpatiocyte: a high-performance simulator for intracellular reaction-diffusion systems

**DOI:** 10.1101/860650

**Authors:** Satya N. V. Arjunan, Atsushi Miyauchi, Kazunari Iwamoto, Koichi Takahashi

## Abstract

**Background:** Studies using quantitative experimental methods have shown that intracellular spatial distribution of molecules plays a central role in many cellular systems. Spatially resolved computer simulations can integrate quantitative data from these experiments to construct physically accurate models of the systems. Although computationally expensive, microscopic resolution reaction-diffusion simulators, such as Spatiocyte can directly capture intracellular effects comprising diffusion-limited reactions and volume exclusion from crowded molecules by explicitly representing individual diffusing molecules in space. To alleviate the steep computational cost typically associated with the simulation of large or crowded intracellular compartments, we present a parallelized Spatiocyte method called pSpatiocyte.

**Results:** The new high-performance method employs unique parallelization schemes on hexagonal close-packed (HCP) lattice to efficiently exploit the resources of common workstations and large distributed memory parallel computers. We introduce a coordinate system for fast accesses to HCP lattice voxels, a parallelized event scheduler, a parallelized Gillespie’s direct-method for unimolecular reactions, and a parallelized event for diffusion and bimolecular reaction processes. We verified the correctness of pSpatiocyte reaction and diffusion processes by comparison to theory. To evaluate the performance of pSpatiocyte, we performed a series of parallelized diffusion runs on the RIKEN K computer. In the case of fine lattice discretization with low voxel occupancy, pSpatiocyte exhibited 74% parallel efficiency and achieved a speedup of 7686 times with 663552 cores compared to the runtime with 64 cores. In the weak scaling performance, pSpatiocyte obtained efficiencies of at least 60% with up to 663552 cores. When executing the Michaelis-Menten benchmark model on an eight-core workstation, pSpatiocyte required 45- and 55-fold shorter runtimes than Smoldyn and the parallel version of ReaDDy, respectively. As a high-performance application example, we study the dual phosphorylation-dephosphorylation cycle of the MAPK system, a typical reaction network motif in cell signaling pathways.

**Conclusions:** pSpatiocyte demonstrates good accuracies, fast runtimes and a significant performance advantage over well-known microscopic particle simulators for large-scale simulations of intracellular reaction-diffusion systems. The source code of pSpatiocyte is available at https://spatiocyte.org.

## BACKGROUND

Intracellular space plays an important role in many biochemical systems operating in the timescales of minutes to hours such as cell signaling [1], division [2], polarization [3], morphogenesis [4] and chemotaxis [5, 6]. These systems are regulated by the spatiotemporal dynamics of molecules. Recent quantitative biology methods can obtain high-resolution spatiotemporal measurements of the molecules. These disparate sources of measurements can be combined and interpreted using spatially resolved simulators to construct models of the systems that are realistic and consistent with physical principles. By simulating the models, detailed analysis of the systems can be conducted in silico and future experiments can be designed [7].

The choice of spatial simulators largely depends on the timescale and spatial resolution of the system of interest [8–13]. For example, high-performance molecular dynamics (MD) simulators [14] can accurately capture the atomistic behavior of up to several macromolecules in a system but are limited in their timescale, allowing only simulations for up to a few milliseconds [15,16]. It is thus not feasible to use MD for example, to simulate cell signaling systems such as the mitogen-activated protein kinase (MAPK) cascade, which takes place at the cellular scale with timescales spanning minutes to hours [17, 18]. For these longer spatial and temporal scales, numerical methods that solve partial differential equations (PDEs) or coarse-grained stochastic particle simulation methods can be used. PDE-based tools such as the freely available Virtual Cell [19] and the commercially available COMSOL Multiphysics (COMSOL Inc.) are useful when the system of interest is deterministic. They are especially fast and convenient when simulating molecules with very high copy numbers. Conversely, particle methods are preferable when we need to account for the noise and fluctuations arising from low copy number of reacting molecules in the cell [20].

Lattice-based particle methods based on the reaction-diffusion master equation (RDME) have the advantage to simulate a large number of diffusing molecules for extended spatiotemporal scales [21–25]. However, since RDME methods represent molecules as dimensionless point particles in lattice voxels, they do not directly capture the effects of excluded volume brought by intracellular macromolecular crowding [26]. About 20–30% of the total volume inside cells are occupied by macromolecules [27]. This amount of crowding has been shown to affect reaction equilibria both in vivo and in vitro, and alter protein binding and gene expression characteristics [28–31]. Moreover, crowded media can also cause non-intuitive effects such as molecules performing directed motion [32], and a change in the statistics of molecular number fluctuations in simple reactions [33]. To capture the effects of volume exclusion in systems that are in equilibrium, Cianci and colleagues [34] recently reported a modified version of the RDME method called vRDME. Despite this enhancement, RDME-based methods are still constrained when it comes to simulating diffusion-limited reactions and rebinding events [35–37] because they assume molecules to be well-mixed in each voxel.

Off-lattice microscopic particle methods such Smoldyn [38], eGFRD [36], SpringSaLaD [39] and ReaDDy [40] can capture the effects of crowding directly because each molecule is represented individually with sphere-like physical dimensions. These simulators also support different sizes of volume excluding molecules. ReaDDy and SpringSalaD can also account for the coarse shape of molecules. However, because of these additional details, microscopic particle methods are more computationally demanding than RDME methods. In a recent performance benchmark of the microscopic methods [41], Smoldyn required the shortest runtime when simulating the well-known Michaelis-Menten reaction-diffusion kinetics.

Spatiocyte [42] is another microscopic method but molecules diffuse on lattice by hopping from one voxel to another. The current stable version of Spatiocyte accounts for volume exclusion by allowing only a single molecule to occupy a voxel at a time. The maximum size of a molecule is roughly equals to the size of a voxel [43]. The method therefore captures steric interactions most realistically when the size of volume excluding molecules is the same and almost equals to that of a voxel. To better simulate the effects of crowding, there is also a development version of Spatiocyte that allows a single molecule to occupy multiple voxels according to its size (unpublished). In Spatiocyte, fine and fast-diffusing molecules such as messengers, metabolites and ions are simulated at the compartment scale using the Next-Reaction method [42, 44]. The accuracy and consistency of Spatiocyte have been validated in detail recently in both volume [43] and surface [45] compartments. Spatiocyte achieves better execution time scaling behavior compared to other methods [43] because it resolves molecular collisions by looking only at the target voxel for occupancy. At the typical intracellular protein concentration range, the performance of Spatiocyte is comparable to Smoldyn when molecules are represented as point particles. On the other hand, when the molecules have physical dimensions, the runtime of Spatiocyte is at least two times faster than Smoldyn. These recent performance benchmarks [41, 43] imply that at present Spatiocyte is one of the fastest methods for simulating individual molecules in crowded systems.

There have been several efforts in the past to improve the performance of particle simulation methods using parallelization approaches. RDME methods have been accelerated on Graphics Processing Units (GPUs) [46–48] and CPU clusters [49]. The GPU-based implementations of RDME can simulate up to two orders of magnitude faster than the serial version on CPU. Chen and De Schutter [49] used Message Passing Interface (MPI) to run a neuron model and achieved 500-fold speedup on a cluster with 1000 processes. Microscopic methods such as ReaDDy [50] and Smoldyn [51, 52] have also been parallelized on GPUs. The performance gain of ReaDDy was up to 115 times over its serial counterpart on CPU. The GPU versions of Smoldyn required between 135- to 200-fold shorter runtimes than the original CPU implementation. Recently, the ReaDDy method was extended to run using multiple threads in parallel on CPU [53]. It showed 6-fold reduction in the simulation time when running with 6 threads compared to the serial implementation.

Here we introduce a parallel implementation of the Spatiocyte method, called pSpatiocyte. The new algorithm was written bottom-up in C++ and MPI to exploit the resources of conventional workstations and massively parallel computers for high-performance simulations of large or crowded cell models. We demonstrate efficient simulations of reaction-diffusion systems at the microscopic scale with volume occupying molecules to recapitulate the crowded nature of intracellular media. We achieve scalability over 500,000 CPU cores on a distributed memory architecture. In the following section we describe the parallelization schemes and numerical implementation of pSpatiocyte. We then provide computational results that validate parallel diffusion and reaction processes. We also demonstrate the performance of pSpatiocyte on the RIKEN K computer with thousands of cores and on a common workstation with eight cores. We show the applicability of pSpatiocyte in actual biological problems by simulating the dual phosphorylation-dephosphorylation cycle of MAPK. Finally, we conclude by providing a summary of the validation and performance results, and future directions of this work.

## METHODS

In a Spatiocyte model, a molecule of a species *s*_*i*_ diffuses in space by performing random walk on lattice from one voxel to a nearest neighbor voxel. The diffusion interval, 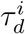 between two successive walks is determined by the diffusion coefficient *D*_*i*_. When the molecule collides with a molecule of a reactant species *s*_*j*_ in the target voxel, they perform a bimolecular reaction with an acceptance probability, *W*_*ij*_. *W*_*ij*_ captures the intrinsic reaction rate *k*_*ij*_ of the pair according to the Smoluchowski-Collins-Kimball (SCK) model [54, 55]. The accuracy and consistency of the bimolecular reaction on lattice in both activation-limited and diffusion-limited regimes have been verified [43, 45]. To represent volume occupying molecules, a voxel can be occupied by a single molecule at any given time. Therefore, if the diffusing molecule meets a non-reactive molecule in the target voxel, a collision occurs and it remains in its original voxel. In the following subsections, we describe the parallelization schemes of the Spatiocyte method in detail.

### Coordinate system

Spatiocyte adopts the hexagonal close-packed (HCP) lattice arrangement (Figure 1A) as it supports the highest density of sphere voxels in a given volume [56]. For comparison, the average density of HCP voxels is 74.048%, whereas the more commonly used cubic lattice has a density of 52.359%. The highest density of voxels is preferable because it allows the simulator to represent highly packed and crowded regions in a compartment to its maximum theoretical limit. Moreover, it was recently demonstrated that the voxels in HCP lattice need to be only about 2% larger than the molecule for the simulations to be consistent with the SCK model [43]. This is in contrast to the cubic lattice, which requires the voxel size to be at least 8% larger. The HCP lattice therefore, can more closely represent hard-sphere molecules in space. For two-dimensional (2D) planar simulations, Spatiocyte employs the triangular lattice arrangement, which is a plane of the HCP lattice [45]. Grima and Schnell [57] have previously shown that simulations on a triangular lattice are closer to Brownian dynamics and produce less discretization error than on square lattice. Simulations on square lattice also overestimate macromolecular crowding effects compared to the triangular lattice.

**Figure 1:**
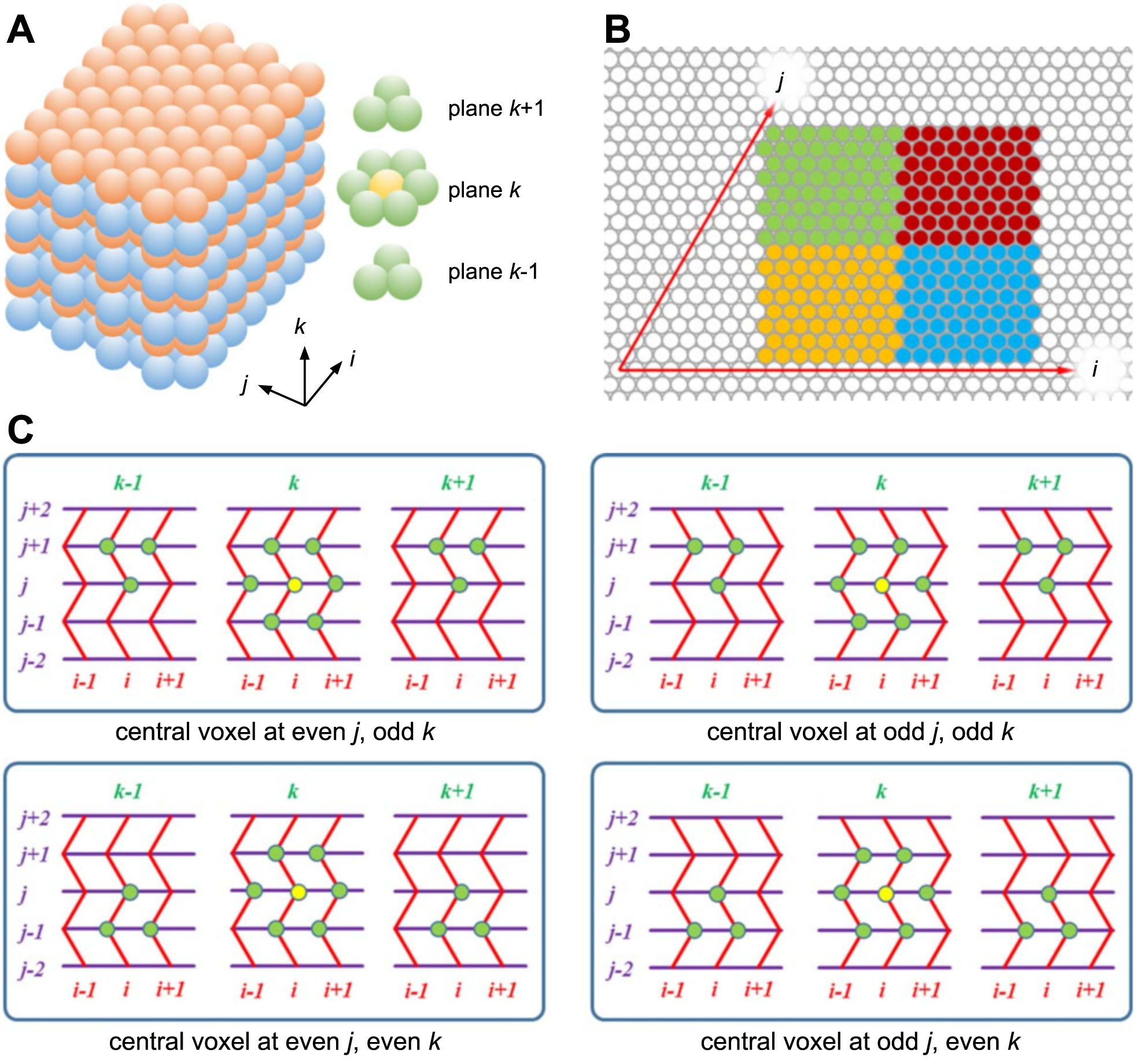
HCP lattice arrangement and a coordinate system for accessing voxels. (**A**) A schematic view of HCP lattice. Left: odd and even planes, shown in different colors. Right: the twelve nearest neighbors (green) of a voxel (yellow). (**B**) 2D slice of an HCP coordinate system based on a unit cell. A rectangular compartment mapped along the axes *i* and *j*, which together make up a parallelogram. A tilted third axis, *k* (not shown) further contorts the boundaries of the compartment. (**C**) Top row: accessing neighbor voxels in a twisted Cartesian coordinate system for the odd plane in *k*-axis. Bottom row: another stencil is used for the even plane.

Although HCP lattice has a regular grid arrangement of voxels, some considerations are necessary to define the coordinate axes to access the voxels. In Figure 1B, the axes *i* and *j* aligned to the voxels in a plane of HCP lattice make up a parallelogram instead of a rectangle. The third axis *k* is also tilted when aligned to the voxels across the planes of the lattice. Such unusual axes arrangement makes it arduous to convert the integer coordinates of a voxel into real coordinates. A data structure without a coordinate system can also be used to identify and access neighbor voxels. For example, a one-dimensional array of voxels can be used, wherein each voxel has pointers to its 12 nearest neighbors. The serial version of Spatiocyte adopts this scheme [42] but from our performance profiling results, the additional memory required to store 12 pointers per voxel and indirect memory accesses to load neighbor voxel data adversely impacts performance because of increased memory bandwidth usage and cache misses. In addition, with an unconventional grid it can also be cumbersome to split the computational domain spatially into smaller subdomains (green, red, orange and blue regions in Figure 1B) for parallel execution by processes.

To overcome these issues, we propose a coordinate system called twisted Cartesian as depicted in Figure 1C. It comprises a straight and two zigzag axes. The figure illustrates how each of the 12 neighbors is identified with these axes. Depending on whether the voxel plane, *k* is even- or odd-numbered, one of two procedures should be used in *i*- and *j*-axes to identify a neighbor voxel. The two procedures are represented by the top and bottom panels of Figure 1C. This coordinate system can be readily mapped onto a conventional Cartesian coordinate system without almost any modification and enables straightforward programmability. Despite being unconventional, the twisted Cartesian coordinate system works well for identifying and accessing neighbor voxels pointer-free by only using conditional statements. In a preliminary implementation of pSpatiocyte, we used the system successfully for parallelization [58]. Recently, a similar approach was adopted for cellular automata simulations in 2D space [59].

#### Algorithm 1: Initialization procedure of the pSpatiocyte algorithm.

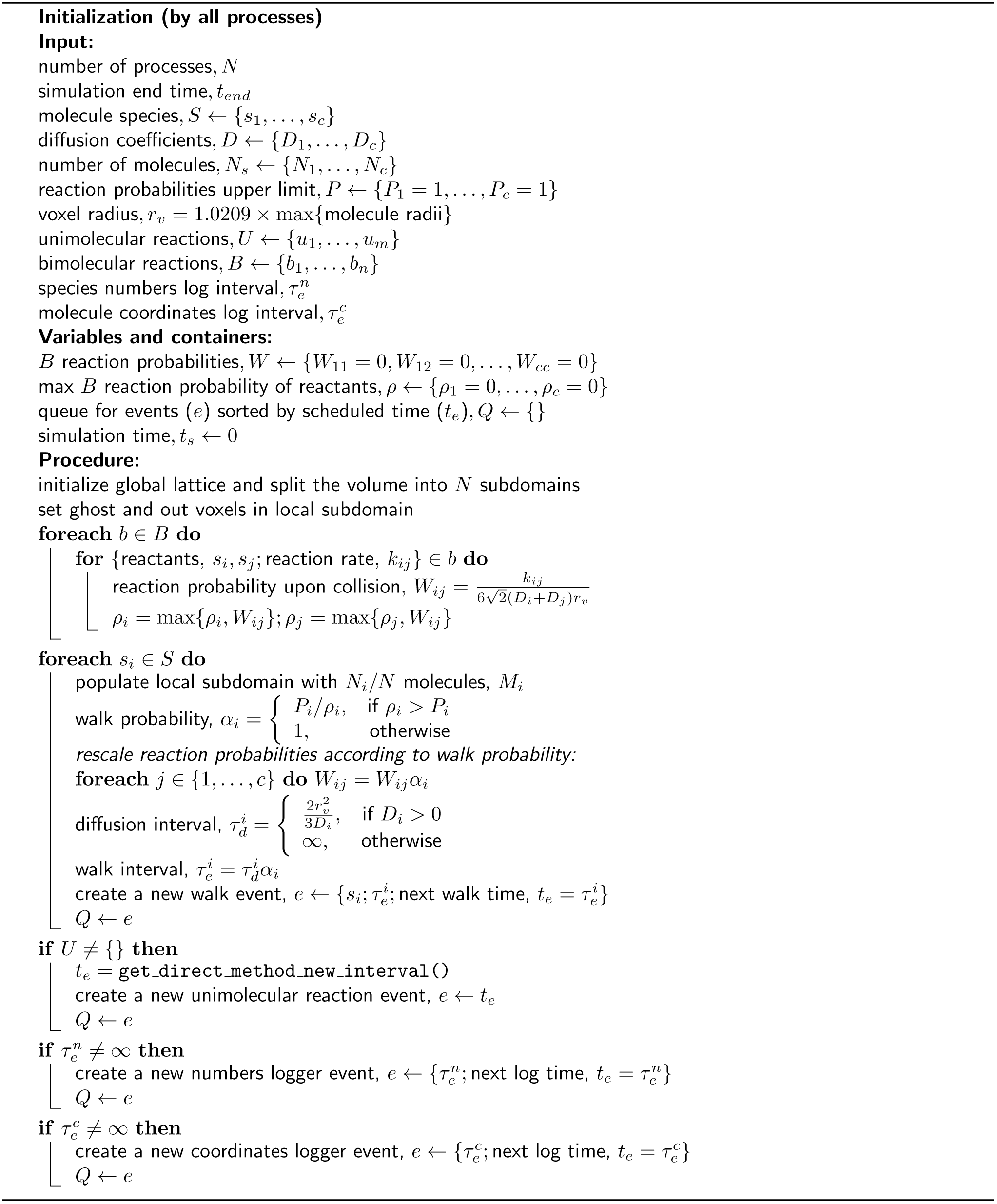

### Parallelized Spatiocyte algorithm

The Spatiocyte method advances the simulation time, *t*_*s*_ in discrete steps using an event scheduler [42,60]. The scheduler gets the next event, *e* to be executed from a priority queue *Q*, which contains events sorted according to scheduled times, *t*_*e*_. The types of events that can be defined in a model include walk (performs diffusion and bimolecular reactions), unimolecular reaction, species numbers logger and molecule coordinates logger. Upon execution, an event returns the interval, *τ*_*e*_ for its next execution. The next execution time, *t*_*e*_ = *t*_*s*_ + *τ*_*e*_ is then passed to the priority queue to reschedule the event. The scheduler executes all events in a loop until the simulation end time, *t*_*end*_ is reached.

The pSpatiocyte method is a parallelized version of Spatiocyte and is completely written in C++. The method is parallelized using the domain decomposition approach illustrated in Figure 2A. With this approach, the complete lattice space is divided equally into *N* subdomains to be executed concurrently by *N* processes. To minimize synchronization overheads between processes, the scheduler and events are duplicated across all processes at initialization, which is described in Algorithm 1. The scheduler of each process then executes the main loop of the simulation in parallel according to Algorithm 2. The simulation proceeds synchronously over all processes by ensuring that (1) the scheduled execution time of each event is identical across processes; and (2) each process synchronizes with adjacent processes when molecules from local subdomain walk or react across adjacent subdomains. We describe how these two conditions are satisfied by each event in the following subsections.

**Figure 2:**
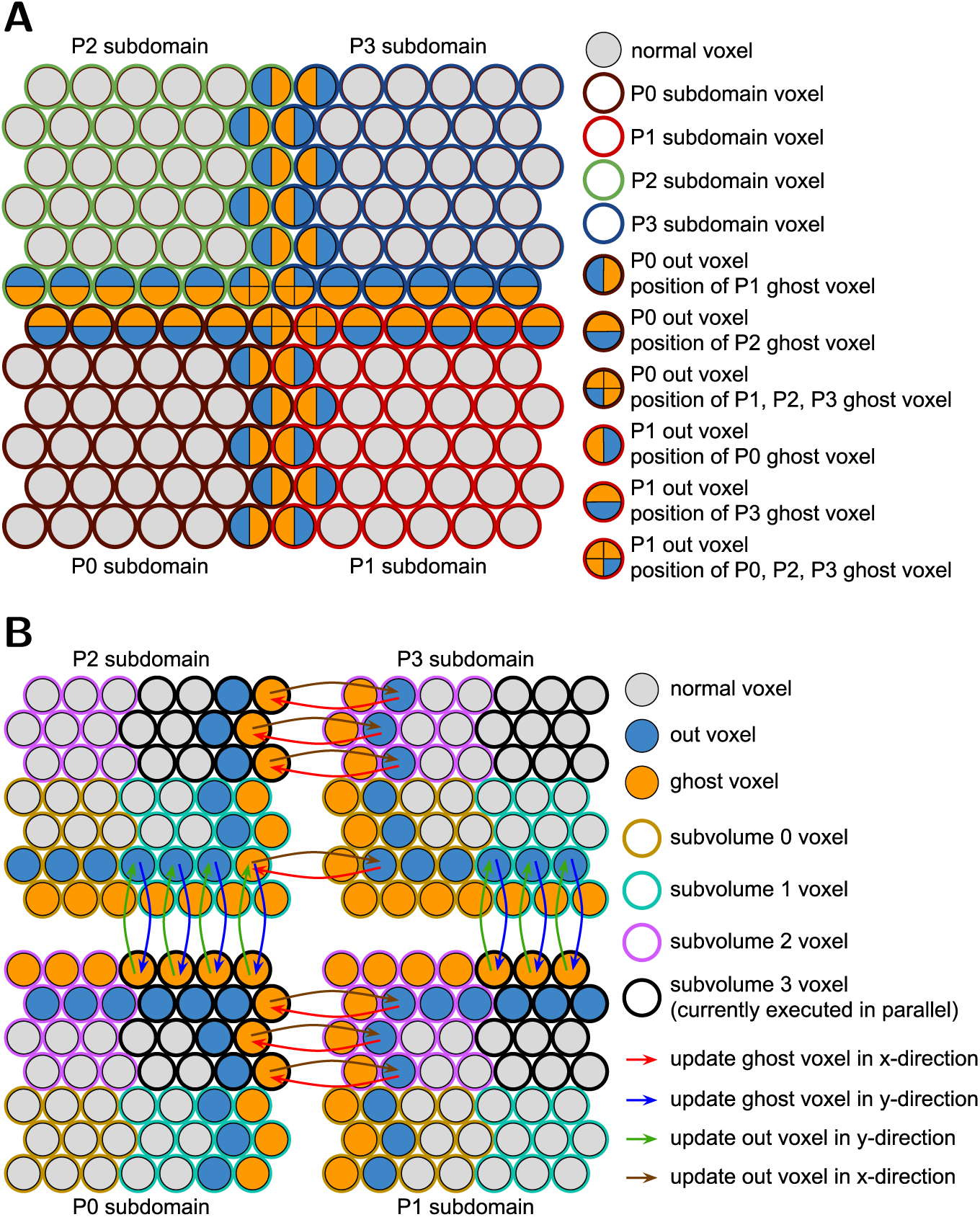
Schemes for large-scale parallel simulation of particles. (**A**) Domain decomposition of an HCP lattice plane. A 12^2^ plane with reflective boundary is equally divided into four subdomains. Each subdomain measuring 6^2^ voxels is allocated to one of four available processes, P0, P1, P2 and P3. Subdomain voxels adjoining other subdomains are defined as out voxels. Ghost voxels are added locally to each subdomain to encapsulate out voxels. The ghost voxels serve to reflect the state of out voxels residing in adjacent subdomains. (**B**) Subdomain division into subvolumes and three-stage inter-process communication. Each subdomain in (**B**) is divided into four equal subvolumes (eight subvolumes, if 3D subdomain). To avoid biased walk events, one of the four subvolumes is randomly chosen before the corresponding local subvolume is executed simultaneously by the four processes. In the above example, subvolume 3 was selected randomly and it is currently being executed in parallel by the four processes. Ghost voxels will be updated using the three-stage communication scheme before they are accessed. The scheme updates the voxels consecutively in x- and y-directions (and z-direction if 3D subvolume). After performing the walk and reaction events in the subvolume, the out voxels in adjacent subdomains will be updated to reflect the state of local subvolume ghost voxels. The updates will be performed successively in (z-,) y- and x-directions. In the example above, the state of an out voxel of P3 subvolume 0 is first transferred to a ghost voxel of P2 subvolume 1 in x-direction before it is communicated to the ghost voxel in P0 subvolume 3 in y-direction. Conversely, the state of the out voxel is updated in reverse, first in y-direction followed by x-direction.

### Parallelized walk event

Einstein [61] and von Smoluchowski [62] have independently shown that small particles in one-dimensional (1D) system perform Brownian walk with a root-mean-squared displacement of 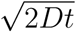, where *t* is the interval between walks and *D* is the diffusion coefficient of the particles. The 1D relation can be expressed in three-dimensional (3D) space with the mean-squared displacement (MSD) given as 6*Dt*. Similarly, in the 3D HCP lattice space, the displacement of a molecule of species *s*_*i*_ within a diffusion interval 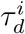 must be consistent with its diffusion coefficient *D*_*i*_. Since the molecule displacement over the interval is equivalent to the voxel diameter, we can use the MSD relation in 3D to obtain the interval, 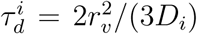, where *r*_*v*_ is the voxel radius. We have previously shown that this approach is accurate for modeling bimolecular reactions on HCP lattice when *r*_*v*_ ≈ 1.0209*R*, where R is the molecule radius [43].

Bimolecular reactions are handled by the walk event because they take place upon the collision of two reactant molecules on lattice during diffusion. Since the reaction acceptance probability, 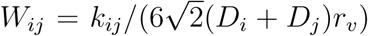 is inversely proportional to the diffusion coefficients of reactants, highly diffusion-limited reactions can cause *W*_*ij*_ > 1 [42, 43]. To address this inaccurate condition, we first determine the maximum *W*_*ij*_ of all bimolecular reactions involving *s*_*i*_ and assign it as *ρ*_*i*_. Then, we obtain the walk probability,

#### Algorithm 2: The main loop of the pSpatiocyte simulation algorithm.

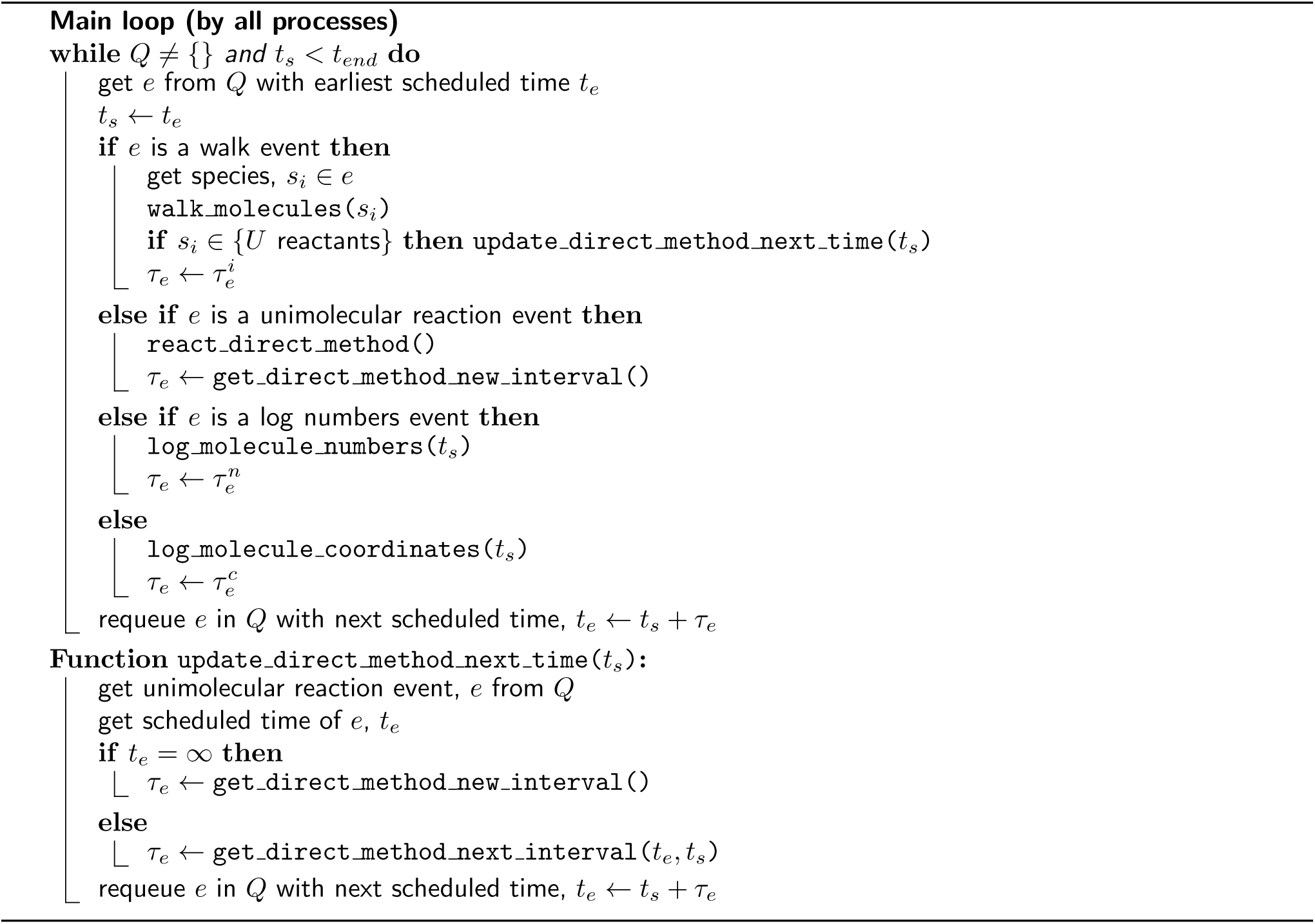

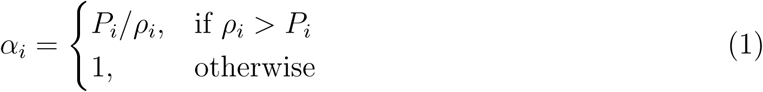

where *P*_*i*_ is a user-defined upper limit of reaction acceptance probabilities and 0 < *P*_*i*_ *≤* 1 (by default, *P*_*i*_ = 1). Next, we rescale the reaction probabilities to *W*_*ij*_ = *W*_*ij*_*α*_*i*_. Finally, we obtain the walk interval as 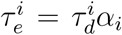. This walk interval is fixed and identical across all processes throughout the simulation procedure. With the above bimolecular reaction scheme, we have shown that the rebinding-time probability distribution of a reactive molecule pair on HCP lattice agrees well with continuum theory (see Section III.A of [43]). We have also verified the accuracy of the reaction rate coefficient and its time-dependent behavior by comparison to SCK theory. It should be noted that reactions that are not highly diffusion-limited have small impact on the simulation performance since they are generally far fewer than diffusion steps (*α*_*i*_ = 1) in the microscopic lattice space.

One of the difficult problems in parallelizing stochastic diffusion and reaction events is maintaining consistency at subdomain boundaries during simulation time steps. Processes should take careful consideration when accessing or writing to voxels residing in adjacent subdomains since they are also simultaneously accessible to the adjacent processes. Figures 2A and 2B illustrate our scheme to achieve consistency during walk and reaction events. We define the voxels at the edge of a subdomain and adjoining other subdomains as *out* voxels. We add a virtual set of voxels called *ghost* voxels locally in each subdomain to represent the current state of out voxels residing in adjacent subdomains. With updated ghost voxels, molecules in a subdomain can walk and react across subdomains seamlessly in a time step without requiring many inter-process synchronization requests. At the end of a walk event, the state of out voxels in adjacent subdomains will be updated to reflect the state of local ghost voxels. Since in a walk event a molecule can at most hop to or react with a molecule in one of its immediate neighbor voxels, only a single layer of ghost voxels is necessary to encapsulate local out voxels (Figure 2B).

To ensure the updated state of ghost voxels remain valid until the end of the walk event, we further divide the subdomain equally into eight subvolumes and execute each subvolume synchronously with all processes. In Figure 2B, four of the subvolumes are shown for each sub-domain. Only the ghost voxels belonging to the selected subvolume is updated before executing the molecules in the subvolume. This scheme ensures out voxels in adjacent subdomains are isolated and free from modification when their corresponding ghost voxels in the local subvolume are accessed.

Algorithm 3 provides the complete pseudocode of the walk and bimolecular reaction procedure in a subdomain. For the walk event, pSpatiocyte uses two random number generators. The first generator is initialized with a seed that is unique to each process, whereas the second generator is initialized with a global seed. With the globally seeded generator, a random number that is drawn locally will be identical for all processes. This scheme reduces communication cost when we need a common random number for all processes. Unless stated otherwise, all random numbers are drawn using the locally seeded generator from a uniform distribution with the interval [0, 1). Both generators use the Mersenne Twister algorithm [63] to generate random numbers.

The walk event executes the random walk of all molecules of a species when it is called. For each molecule *m* of species *s*_*i*_, a random target voxel, *v*_1_ out of 12 neighbor voxels is first selected. If *v*_1_ is a ghost voxel, *m* is appended to a list 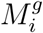, containing molecules targeting ghost voxel, while *v*_1_ is added to *V*_*g*_, a list of the targeted ghost voxels. However, if *v*_1_ is not a ghost voxel and is vacant, a random number *r* is drawn. If *r* is less than or equals to the species walk probability *α*_*i*_, the walk is successful and *m* is moved to *v*_1_. Otherwise, if *v*_1_ contains a reactant pair of species *s*_*j*_, then a random number *r* is drawn. If *r* is less than or equals to the bimolecular reaction probability *W*_*ij*_, then the reaction is performed. If *v*_1_ is instead occupied by non-reactive molecule, a collision occurs and *m* stays in its current voxel.

#### Algorithm 3: The walk event function for species *s*_*i*_.

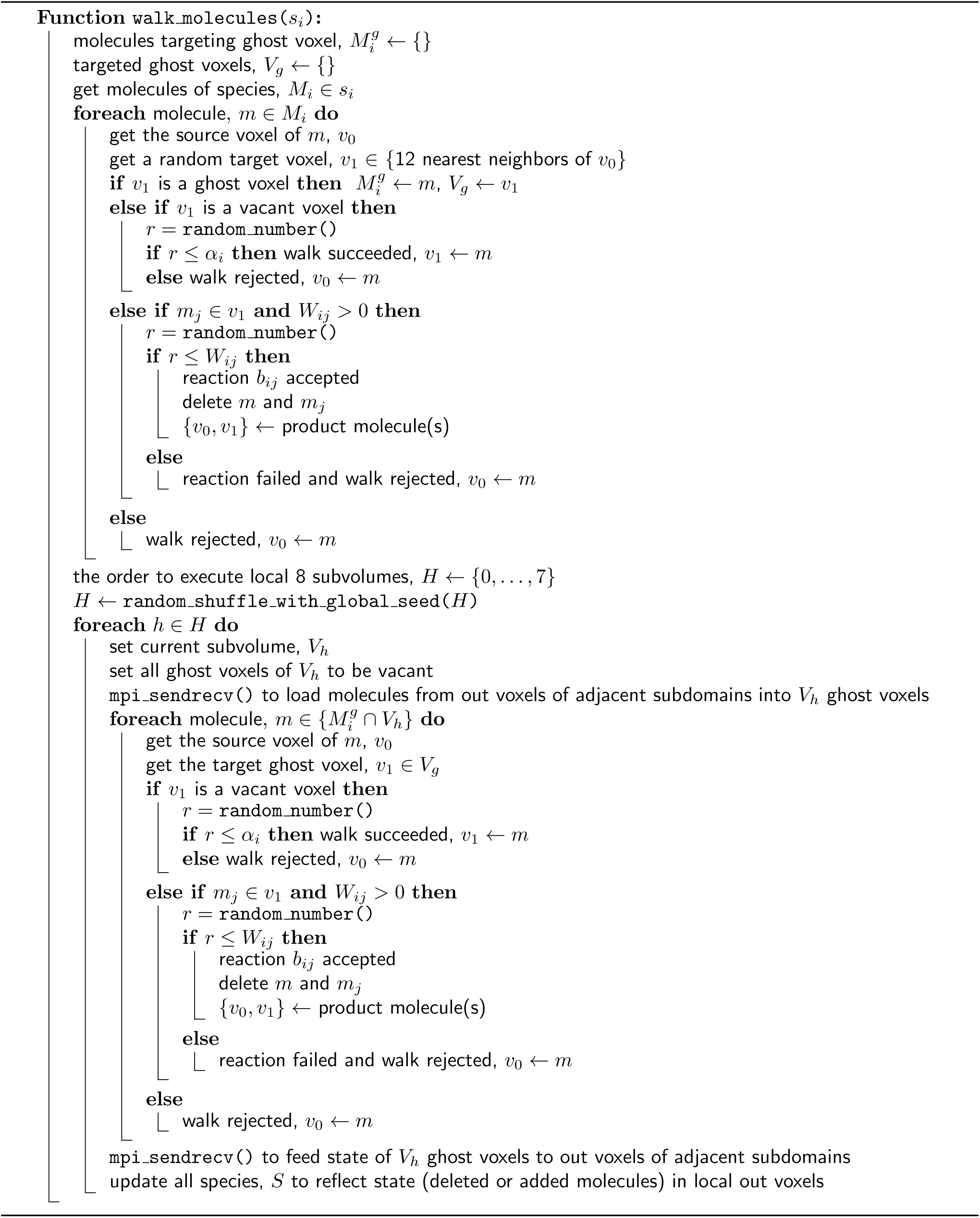

After completing the above procedure for all molecules of *s*_*i*_, a list containing the execution order of eight subvolumes is randomly shuffled using the globally seeded random number generator. For each subvolume *V*_*h*_ in the ordered list, we first update its ghost voxels by loading the state of corresponding out voxels from adjacent subdomains. The update is performed using the MPI Sendrecv function. Next, for each molecule *m* in 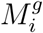 that is in the current subvolume *V*_*h*_, we get its target ghost voxel *v*_1_ from *V*_*g*_. If *v*_1_ is vacant, a random number *r* is drawn. If *r* is less than or equals to the species walk probability, *α*_*i*_ then the molecule is moved to the target voxel *v*_1_. Otherwise, the walk fails and the molecule stays in its voxel. If *v*_1_ is instead occupied by a reactive pair of species *s*_*j*_, then a random number *r* is drawn. If *r* is less than or equals to reaction acceptance probability *W*_*ij*_, the reaction is executed. Otherwise, the reaction fails and the molecule stays in its voxel. After executing all the molecules in the subvolume, we update the out voxels in adjacent subdomains with the state of their corresponding ghost voxels using MPI Sendrecv function. The process then repeats the above procedure with the next subvolume in the ordered list and continues until all subvolumes have been executed.

Note that the walk event in each subvolume is performed locally at all times by each process. Although mutual exclusion is naturally realized by this scheme, inter-process communication is performed once at the beginning and again at the end of each subvolume execution. This scheme of mutual exclusion however does not necessarily require eightfold communication requests because the number of voxels to be sent or received is also reduced in proportion to the subvolume boundary surface area. We have confirmed the effectiveness of this method with at least a few thousand processes [58].

Another problem common in lattice-based parallel computations is the communication at the subdomain vertices. Out voxels located at the vertices are accessed by many more processes than at other locations. Generally, latency has the most impact during the short communications required at these voxels. In addition, contention between requests tends to occur because of the limited bandwidth or the number of available communication channels. From the viewpoint of strong scaling, the communications will show poor performance especially when involving large number of processes. To overcome these constraints, we employed the three-stage communication scheme, which is well-established and described previously [64]. With this scheme, it is sufficient for each process to communicate with six directly adjacent processes in three consecutive stages. In our implementation, we adapted the scheme to update ghost and out voxels when executing each of the eight subvolumes. An example of the communication scheme is illustrated in Figure 2B.

### Parallelized unimolecular reaction event

The sequential Spatiocyte method employs the Next-Reaction method for unimolecular reaction events [42, 44]. However, in the pSpatiocyte method, we have adopted the Gillespie’s direct-method [65] to execute the events in parallel because of its simplicity. In the direct-method, the propensity for a unimolecular reaction, 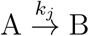 is *a*_*j*_ = *k*_*j*_*X*_*A*_, where *X*_*A*_ is the number of A molecules in the volume. If there are *m* unimolecular reaction channels, the total propensity is given by

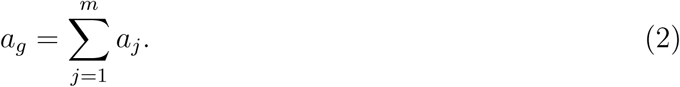

#### Algorithm 4: Parallelized Gillespie’s direct-method for unimolecular reactions.

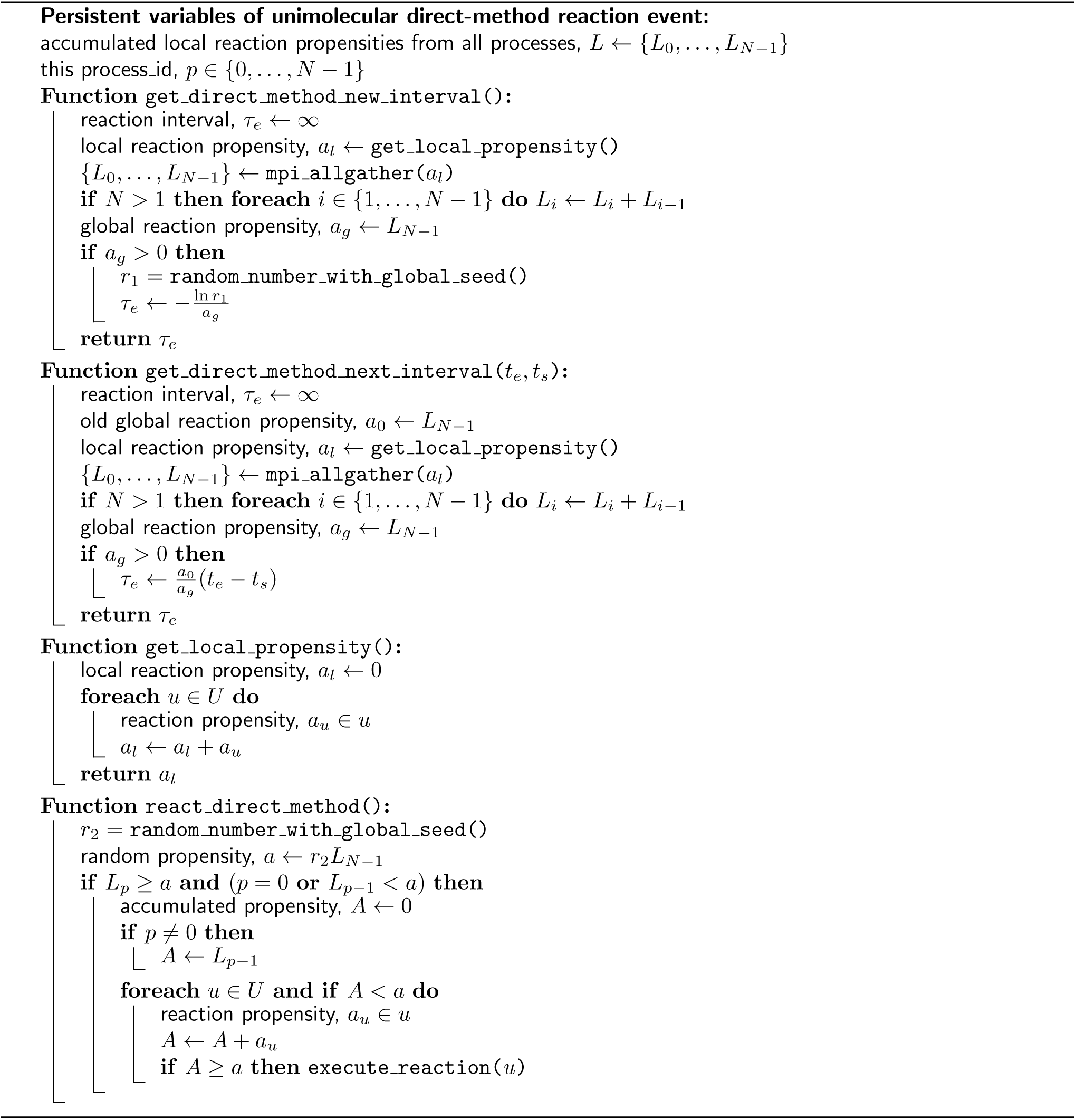

The interval, *τ*_*e*_ for the next reaction event is expressed as

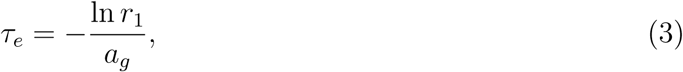

where *r*_1_ is a random number drawn from a uniform distribution in the unit interval. At the end of the interval, the reaction channel *u*, out of the total *m* channels is selected to be executed such that

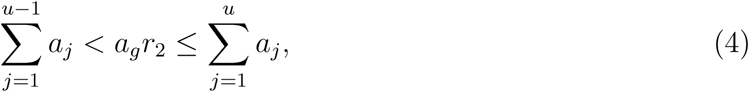

where *r*_2_ is another uniform random number from the unit interval.

We have parallelized the direct-method using two simple schemes. First, in each process, we use MPI Allgather to get the local propensities, *a*_*l*_ from all subdomains and sum it locally to get the global propensity, *a*_*g*_. Second, we use the globally seeded random number generator to draw *r*_1_ and *r*_2_ to determine *τ*_*e*_ and *u*, respectively. With these two schemes, *τ*_*e*_ and *u* will be identical for all processes without additional synchronization requests. The pseudocode of the parallelized direct-method is given in Algorithm 4.

At initialization, the scheduler gets the next time to execute the unimolecular reaction event, *t*_*e*_ = *τ*_*e*_ by calling the `get_direct_method_new_interval` function and schedules it in the priority queue, *Q*. The new interval is calculated from Eq. (3). In the main loop of the simulation, if a walk event of a unimolecular reactant species has been called, the number of reactant molecules may have changed. Therefore, at the end of the walk event, the next time of the reaction event is updated by calling the `get_direct_method_next_interval` function. It gets the updated interval from the scaling expression, *τ*_*e*_ = *a*_0_(*t*_*e*_ − *t*_*s*_)*/a*_*g*_, where *a*_0_ and *a*_*g*_ are the old and new global propensities, respectively, *t*_*e*_ is the old scheduled time and *t*_*s*_ is the the current simulation time. Finally, the scheduler executes the reaction event at the scheduled time by calling the `react_direct_method` function and reschedules it using a new interval from `get_direct_method_new-interval`. The `react_direct_method` function selects the reaction channel to be executed according to (4).

When the reaction channel is executed, if there are two product molecules, one of them will replace the reactant in its current voxel. Another random vacant voxel from the 12 nearest neighbors of the reactant will be selected to occupy the second product. When the compartment or the region near the reactant is highly crowded, no vacant voxel may be found for the product. In the original Spatiocyte method, this will result in a failed unimolecular reaction. We have also adopted this approach for the pSpatiocyte method. In addition, for such a highly crowded scenario, we have added an option in the model to randomly vacate one of the neighbor voxels of the reactant and place the second product in it. The voxel can only be vacated if the molecule occupying it is a mobile (diffusing) species. The voxel is first vacated by moving the molecule to a vacant neighbor voxel. If no vacant voxels are available for the molecule, the procedure is repeated with another randomly selected neighbor molecule of the reactant. The reaction fails if none of the nearest neighbors of the reactant can be vacated for the second product.

### Parallelized logger events

Two types of logger events are available to save the snapshots of pSpatiocyte simulation. The species numbers logger event saves the number of molecules of each species in a comma-separated values (CSV) file when called by the scheduler. At initialization, the log interval is fixed and duplicated across all processes, ensuring that the scheduled execution time of the logger event is always identical across processes. During simulation, each process writes the molecule numbers available in its subdomain into a local file, thus avoiding inter-process communication. A Python script is provided to gather and sum all the numbers from the process files into a single conventional CSV file after the simulation.

The coordinates logger is implemented the same way as the numbers logger but it saves the unique identity number (ID) and integer (voxel) coordinates of each molecule within its subdomain. The ID of a molecule is persistent across subdomains, hence it is possible to track the trajectory of each molecule throughout the simulation. The logger can also save the coordinates of out and ghost voxels for debugging purposes. A Python script gathers and converts the integer coordinates into real 3D coordinates before saving them into a CSV file.

## RESULTS AND DISCUSSION

To confirm the consistency of pSpatiocyte in terms of physical accuracy, we validated diffusion and reaction processes when running on all eight cores of a workstation with Intel Core i9-9900K CPU (8 cores, 5 GHz maximum processor frequency), 64 GB memory and Ubuntu 19.10 operating system. Parallel performances of diffusion were examined across thousands of computational cores of the K computer [66]. pSpatiocyte performance was also compared with Spatiocyte, Smoldyn and ReaDDy when simulating the benchmark enzymatic reaction model on the workstation. Finally, the parallelized simulation outcomes of the well known MAPK model are provided as an application example. All simulations were performed on lattices with reflective boundaries.

### Validation of diffusion

We initially observed the trajectories of molecules diffusing across subdomains to verify the coordinates logger, inter-process communications and the overall simulation algorithm. Figure 3A displays the trajectories of five molecules diffusing with *D* = 0.06 *µ*m^2^s^−1^ for 10 s. The simulation was executed with eight processes, employing all cores of the workstation. At the beginning of the simulation, the molecules were placed randomly in a compartment volume of 10 *µ*m^3^ with lattice dimensions 476^3^, divided into eight subdomains. All trajectories in Figure 3A appear consistent with molecules performing Brownian motion.

**Figure 3:**
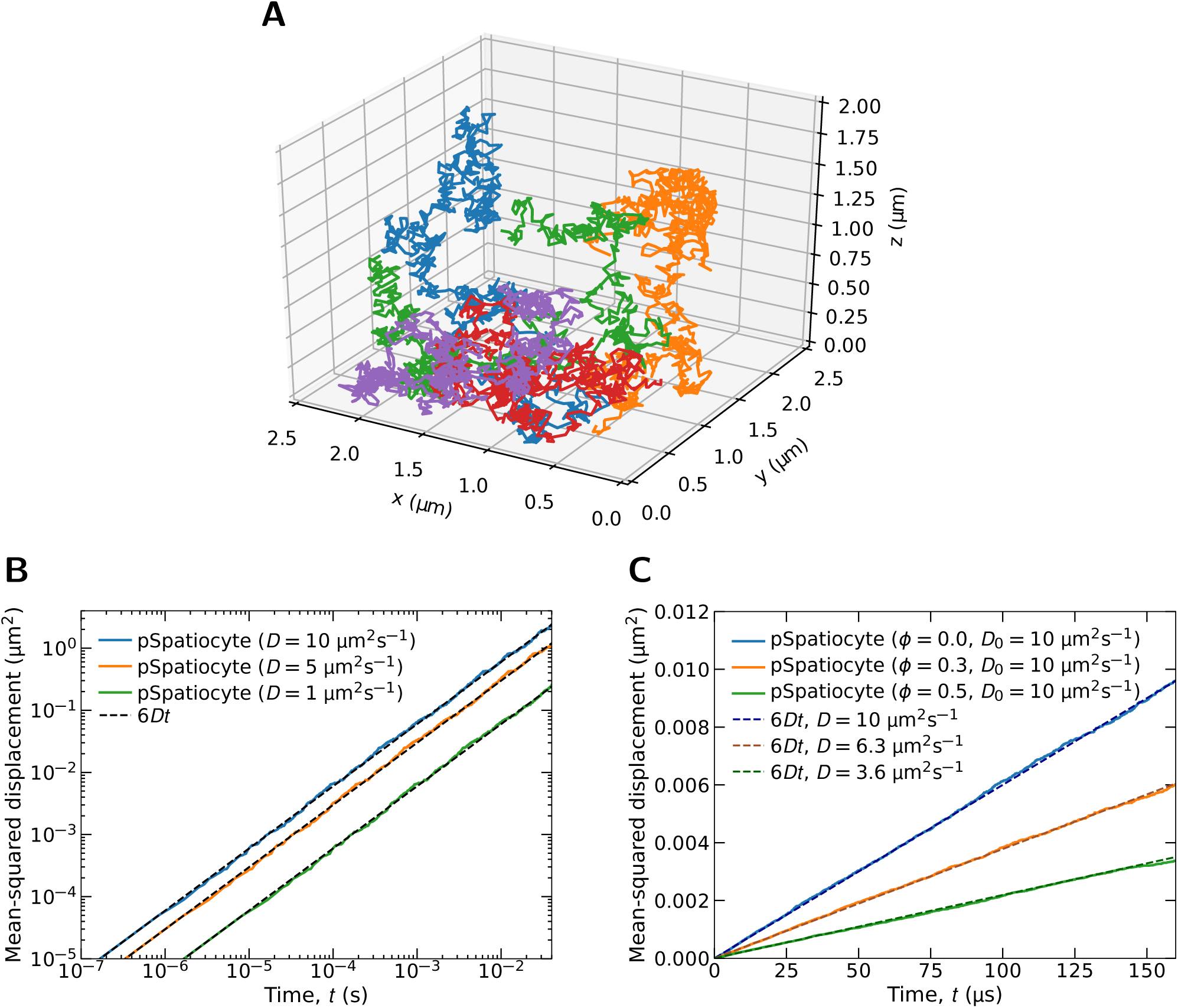
Parallelized 3D diffusion in dilute and crowded conditions. (**A**) Trajectories of five molecules diffusing across lattice subdomains. (**B**) Mean-squared displacements (MSDs) of molecules diffusing in a sparsely populated compartment. (**C**) MSDs of molecules diffusing in crowded conditions. *D*_0_ is the diffusion coefficient specified in the model, whereas *D* is effective rate fitted to the resulting MSD. The fraction of voxels occupied by immobile crowder molecules are indicated by *ϕ*.

We then validated the consistency of pSpatiocyte in reproducing the correct diffusion behavior in a dilute volume that is equally distributed to eight processes. The MSD of a molecule performing random walk in 3D space is given as 6*Dt*. We monitored the diffusion of a single molecule placed at the center of a 960^3^ lattice with voxels measuring 2.5 nm in radius. No other molecules were present on the lattice. We then performed random walks repeatedly with the same initial conditions aside from the random number generator seed. 100 random walks were performed for 40 ms and the average MSDs were computed from the ensembles. Figure 3B shows the log-log plots of the results for three different diffusion coefficients. The slopes, the vertical distances, and the absolute values coincide well with the expected theoretical lines.

When a compartment is crowded with obstacles as in the cell [67], the effective rate of diffusion is expected to decrease. To evaluate if pSpatiocyte is able to replicate the rate reduction when running with eight processes, we obtained the MSD of a diffusing molecule in a compartment occupied by immobile crowder molecules. The fraction of compartment voxels occupied by the crowder molecules is given by *ϕ*. We evaluated three different *ϕ* conditions, with 1000 independent simulation runs for each condition. Each run adopted a unique seed for drawing random numbers. Hence, the random placement of immobile crowders at initialization was different in each run. The averaged MSDs and the fitted effective diffusion rates are displayed in Figure 3C. As expected, a significant decrease in the diffusion rate corresponding to an increase in *ϕ* is clearly observed. Further detailed analysis is required to compare the effective diffusion rates in crowded condition on HCP lattice with the rates in continuous space as reported by Novak et al. [68].

### Validation of reactions

We validated parallelized irreversible and reversible reactions separately because they have distinct underlying physics.

#### Irreversible reaction

A unimolecular reaction given by

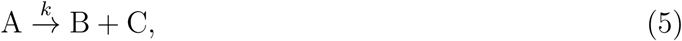

is irreversible. In pSpatiocyte, the reaction is executed according to the parallelized Gillespie’s direct-method. We applied three different reaction rates, *k* in an uncrowded volume and compared the results with that of an ordinary differential equation (ODE) solver. The simulation was executed on eight CPU cores with parameters *D* = 10 *µ*m^2^s^−1^, voxel radius *r*_*v*_ = 5 nm and 960^3^ lattice size. The initial number of reactant molecules was 64,000. Figure 4A and 4B show the simulation results of the reactant and products, respectively. In all cases, the outcomes of simulation agree very well with the ODE solver.

**Figure 4:**
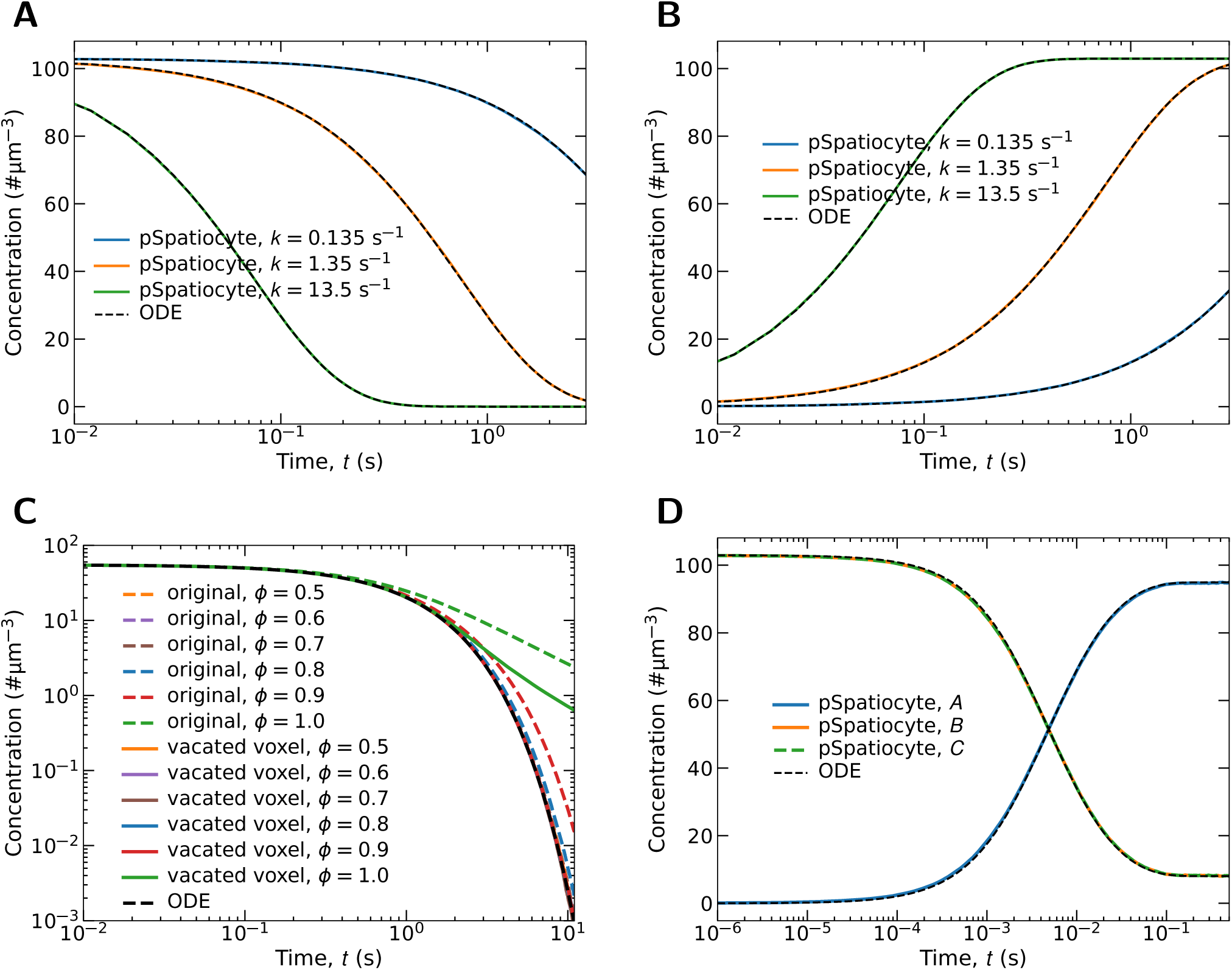
Parallelized unimolecular and bimolecular reactions. (**A**) Time profiles of A in the dissociation reaction 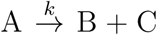. (**B**) The corresponding time profiles of B and C. (**C**) Effects of volume occupancy on the rate of the dissociation reaction on HCP lattice. Curves indicate the concentration profile of *A. ϕ* denotes the fraction of voxels occupied by crowder molecules. In the original method, the reaction fails if there are no vacant voxels among the 12 nearest neighbors of the reactant molecule to place the second product molecule. In the vacated voxels approach, a diffusing molecule from a nearest neighbor is selected randomly and moved to one of its nearest neighbors to allocate a vacant voxel for the product. Simulation model parameters: total volume was 90 *µ*m^3^ with 64^3^ lattice, initial number of A molecules was 50,000, crowder molecules were added to achieve *ϕ* as shown, the diffusion coefficient of A, B, C and crowder molecules was 10 *µ*m^3^s^−1^, and 100 runs for each *ϕ*. (**D**) Time profiles of reactants and products in the reversible reaction 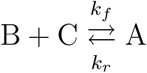.

We investigated the effects of crowding on the dissociation rate of (5), with the two different ways of finding a vacant voxel for the second product molecule. In the original approach, the reaction fails if all neighbor voxels of the reactant are occupied. In the second approach, if they are all occupied, the simulator attempts to vacate one of them. The reaction also fails if no voxels can be vacated for the product. Figure 4C shows the log-log plots of the reactant concentration using the two approaches when *ϕ* is between 0.5 and 1.0. At *ϕ* = 1.0, all voxels of the HCP lattice are occupied, giving about 74% volume occupancy. With the original approach, the dissociation rates agree well with the ODE result up to *ϕ* = 0.7, which translates to about 52% volume occupancy. In the second approach, the rates are comparable up to *ϕ* = 0.9. In vitro results of crowding experiments showed that the dissociation rate of molecules are unaffected even when the volume occupancy reaches 30% [69]. However, it is still unknown if occupancies above 50% would affect the dissociation rate as we have found with our original approach. The results show that the original approach is sufficient for simulating the estimated 30% volume exclusion in the cytoplasm [70]. In cases where the dissociating molecules are much smaller that the voxel size, such as messengers, metabolites and ions, the sequential version of Spatiocyte simulates them at the compartment scale using the Next-Reaction method. Since this feature is not yet supported by pSpatiocyte, we leave it for future work

#### Reversible reaction

A bimolecular reaction is dependent on the diffusion of reactant molecules because the reaction takes place as the molecules meet and collide in space. For simplicity, we considered a forward bimolecular reaction with a single product and a reverse unimolecular reaction, as an example of reversible reaction,

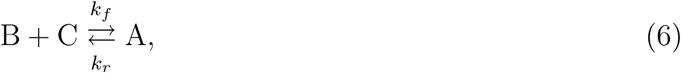

where *k*_*f*_ and *k*_*r*_ denote the effective forward and reverse reaction rates, respectively. The effective forward rate can be converted to the intrinsic rate that is used by pSpatiocyte,

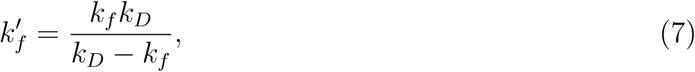

where *k*_*D*_ = 8*πr*_*v*_(*D*_*B*_ + *D*_*C*_) [43]. *D*_*A*_ and *D*_*B*_ are the diffusion coefficients of B and C, respectively. The intrinsic reverse reaction rate is given as

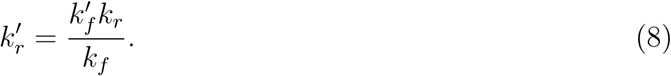

The forward reaction is executed by the walk event when molecules of the reactant species collide on lattice, whereas the reverse reaction is performed by the unimolecular reaction event. The parameters of the simulation include *k*_*f*_ = 2 *µ*m^3^*s*^−1^, *k*_*r*_ = 1.35 s^−1^, *D* = 10 *µ*m^2^s^−1^ and *r*_*v*_ = 5 nm. The initial number of A and B molecules was 64,000 each and the lattice size was 960^3^. The simulation volume was distributed to eight processes. As comparison, an ODE solver was used to generate the output of the reversible reaction with the effective rates, *k*_*f*_ and *k*_*r*_. The results of the simulation are provided in Figure 4D. The closely matching curves of pSpatiocyte and the ODE solver verify the parallel simulation accuracy of reversible bimolecular reactions.

### Performance of parallelized 3D diffusion

In Spatiocyte simulations, diffusion of molecules typically takes place at step intervals that are several orders of magnitude smaller than that of reactions if they are not highly diffusion-limited. The fine intervals are needed for the very short displacements between voxels. Since diffusion computations at these fine intervals dominate the total computation cost of most simulations, we used a diffusion model to evaluate the parallel simulation performance of pSpatiocyte. The simulation parameters and conditions are the same as in the diffusion model in the previous section, except for the lattice resolution and occupancy.

To estimate the parallel performance of pSpatiocyte, we measured its strong and weak scaling efficiencies. Strong scaling measures how fast a program is able to process a fixed workload by splitting it into smaller sizes and distributing them to an increasing number of CPUs or cores. On the other hand, weak scaling measures how large of a problem a program can handle without the loss of speed. To measure the efficiency of weak scaling, the amount of workload given to each CPU or core is fixed and the total workload processed by the program is increased by adding CPUs or cores with the corresponding amount of workload.

For strong scaling, we used three different voxel radii to measure performance. Given that the physical dimensions of a compartment remain the same, smaller voxels would result in higher resolution and finer lattice. We denote voxels having 10, 5, and 2.5 nm radii as coarse, intermediate, and fine lattices, respectively. The molecule occupancy, *ϕ* was fixed at 0.3, whereas the voxel sizes determined the spatial resolution of the compartment. The compartment resolution was 512^3^ (coarse), 1024^3^ (intermediate) or 2048^3^ (fine). For computations using the maximum 663552 cores of the K computer, the resolution was set to 512 × 480 × 540 (coarse), 1024 × 960 ×1080 (intermediate) or 2048 × 1920 × 2160 (fine) to ensure that the model conforms to the physical configuration of the processes.

Note that we examined the relative speedups instead of the floating point operations per second (FLOPS) because on lattice, integer or logical instructions dominate the overall computational cost. The speedups measured from the elapsed times are shown in Figure 5A. Here, we used the results of the coarse lattice with 64 cores as the baseline reference to calculate the speedups since simulations with fewer cores were not possible due to memory size limitation. For the intermediate and fine lattices, the results from 64 cores were extrapolated using the time consumed per voxel on the coarse lattice. The parallel efficiencies of strong scaling are summarized in Figure 5B. For the fine lattice with 663552 cores, we obtained a speedup of 7686. The corresponding strong scaling efficiency was at 74.1%. In contrast to the coarse and intermediate lattices, the results from the fine lattice were the closest to the ideal curve. By extrapolating the results from Figure 5A, we can predict that the speedup and efficiency on the fine lattice would be about 13000 times and at 40%, respectively if two million cores were utilized.

**Figure 5:**
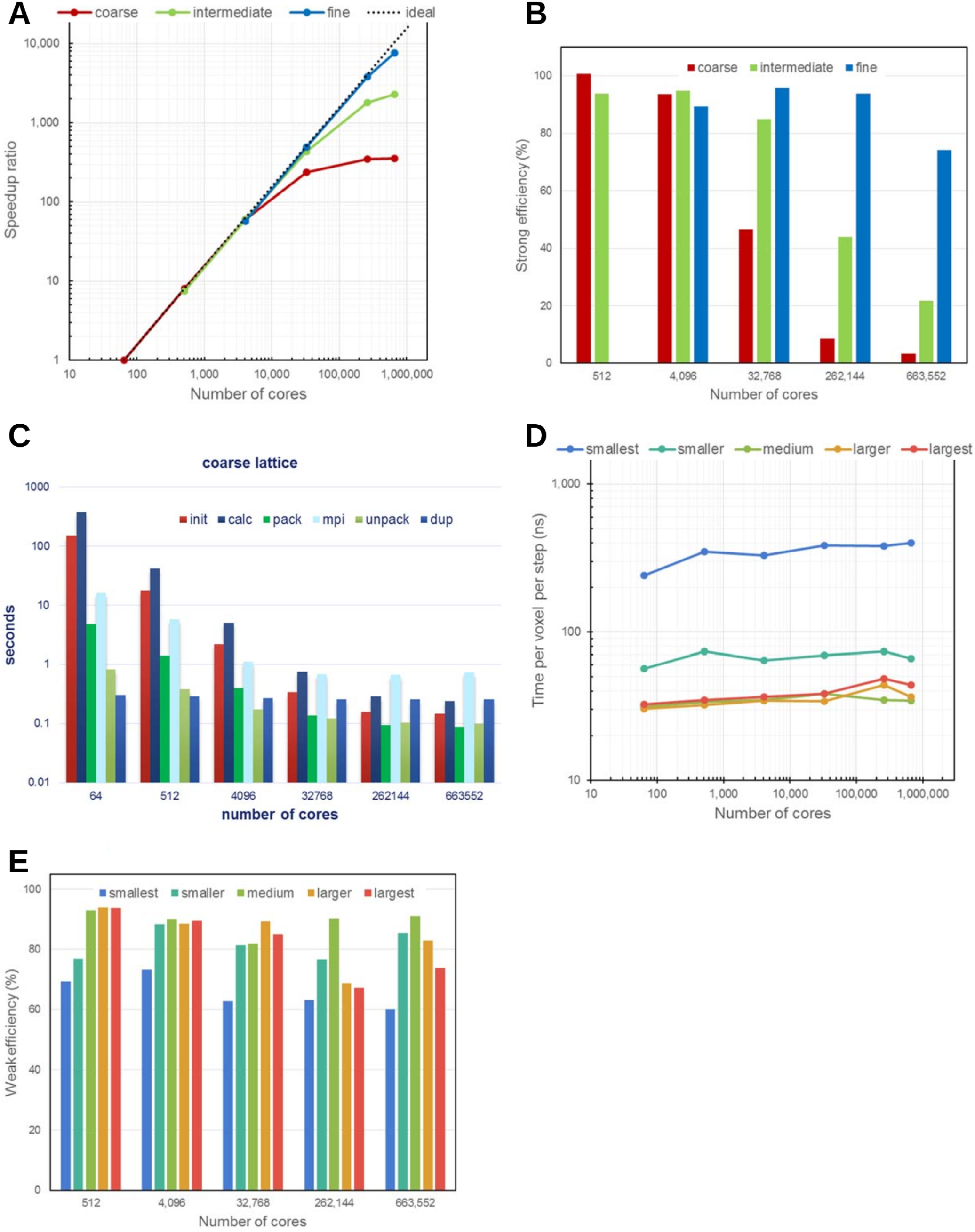
Parallelized 3D diffusion performance of pSpatiocyte. Diffusion model is the same as in Figure 3 but with 30% of the total voxels occupied by diffusing molecules. (**A**) Speedup ratios relative to 64 cores. Red, green, and blue lines represent lattices with 512^3^ (coarse), 1024^3^ (intermediate), and 2048^3^ (fine) voxels, respectively. (**B**) Efficiency of strong scaling on increasing number of CPU cores. The resolutions of lattice are the same as in **A**. (**C**) Components of simulation elapsed times. The average times taken for initialization and computation are given by `init` and `calc`, respectively. For inter-process communications, the `pack` and `unpack` times are the durations required to manage data before sending and after receiving it, respectively. The time spent on the `MPI Sendrecv` function is denoted by `mpi`. The duration to duplicate communicator objects is indicated by `dup`. (**D**) Average elapsed times per voxel per simulation step. Voxel sizes from `smallest` to `largest` denote 32^3^, 64^3^, 128^3^, 256^3^, and 512^3^ voxels per process, respectively. (**E**) Efficiency of weak scaling on increasing number of CPU cores. Colors have the same representation as in **D**.

To identify the cause of the performance deterioration on the coarse lattice, we determined the major components of elapsed time as shown in Figure 5C. We found that the times taken for initialization, computation, pack and unpack events were decreasing at least with up to 262144 cores, whereas the MPI time saturated and exceeded these times when it was over 32768 cores. To improve the performance, the constant duplication time due to redundant communicator objects should be eliminated by sophisticated programming. The saturation of MPI time is likely the most significant factor that needs to be addressed to improve the scaling performance further. However, such saturated timings generally originate from the latency of inter-process communication, which is dependent on the hardware and firmware. Therefore, the immediate approach for improving strong scaling is to reduce the computation time and ensure that it is as close to the latency as possible.

Figure 5D displays the weak scaling performance of pSpatiocyte in elapsed time per voxel per step. Here, the labels on the horizontal axis, smallest, smaller, medium, larger and largest, denote the subdomain lattice dimensions, 16^3^, 32^3^, 64^3^, 128^3^ or 256^3^ on each process, respectively. In an ideal weak scaling performance setting, these times would be identical for all lattice sizes since they would be independent of the number of cores and voxel sizes. In spite of the variation in the absolute values, all lattice dimensions provided good scaling properties. We scrutinized the elapsed times of the smallest and smaller lattices and it revealed that the times are dominated by the constant communication latency. This explains the source of the larger absolute times in the smaller lattices. The efficiencies of parallel computation in terms of weak scaling are summarized in Figure 5E. For each lattice size, the elapsed time with 64 cores was used as a reference to calculate the parallel efficiency. Although the efficiencies tend to deteriorate on higher number of cores, we were still able to achieve 60% efficiency with more than half a million cores.

### Performance benchmark

We compared the runtime of pSpatiocyte with Spatiocyte (git 5e88f40), Smoldyn (v2.61) and ReaDDy (v2.0.2-py37 55 g78bd07) when executing the benchmark Michaelis-Menten enzymatic reaction model [41] on the same workstation. ReaDDy has both serial and parallel versions. All simulators were evaluated on a single core. Additionally, the parallel simulators pSpatiocyte and ReaDDy were executed on two, four and eight cores. We used the same model parameters as in [41,71] but increased the size of the reaction volume tenfold to 909 *µ*m^3^. The larger volume raises the computational cost to an adequate level when running the model in parallel on all eight cores of the workstation. For Smoldyn and ReaDDy, we set the simulation interval, Δ*t* to 1 ms. In pSpatiocyte and Spatiocyte models, the event with the smallest interval is the walk event and it was set to 0.5 ms. We also evaluated pSpatiocyte with a smaller walk event interval of 0.2 ms to see how the runtime scales with the interval. Smaller simulation interval results in an overall higher computational cost and up to a certain extent, better accuracy.

Figure 6A displays the results of the performance benchmark. The runtimes shown are the averages of three independent runs of each simulator to execute the model for 10 s. The resulting concentration profiles from each simulator are plotted in Figure 6B. On a single core, pSpatiocyte with Δ*t* = 0.5 ms is about two times faster than with Δ*t* = 0.2 ms. It is also about 2.4 times faster than the sequential version of Spatiocyte. On a single core, the main difference between pSpatiocyte and its sequential counterpart is the new pointer-free voxel accessing scheme adopted by the former. This scheme has likely lowered the memory bandwidth usage and cache misses, contributing to the significant reduction in simulation runtime. The execution time of pSpatiocyte is about 7.7 times shorter than Smoldyn on a single core. It is also roughly 30- and 38-fold times faster than the serial and parallel versions of ReaDDy, respectively.

**Figure 6:**
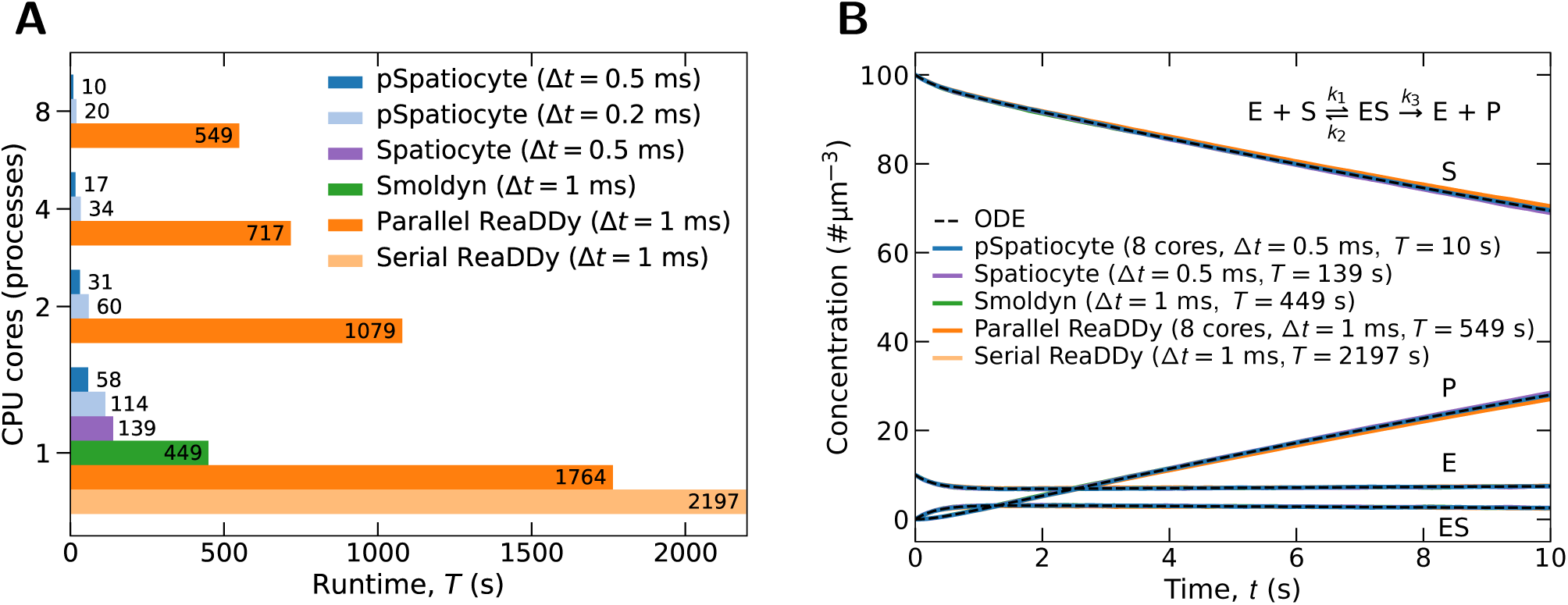
Performance benchmark of the Michaelis-Menten reaction on a workstation. Model parameters [41, 71]: total volume 909 *µ*m^3^, diffusion coefficient 10 *µ*m^2^s^−1^, *k*_1_ = 0.01 *µ*m^3^s^−1^, *k*_2_ = *k*_3_ = 1 s^−1^. The initial numbers of E and S molecules are 9090 and 90910, respectively. Duration of simulation 10 s. Simulation or walk event interval (Δ*t*) and runtime (*T*) are as indicated. (**A**) Comparison of simulator runtimes on different number of CPU cores. (**B**) Concentration profile outcomes from the simulators.

The runtime of pSpatiocyte (Δ*t* = 0.5 ms) also scales favorably with the number of additional cores used in the simulation. When the number of cores was increased from one to two, the runtime was reduced by 1.87 times. Similary, the runtime was shorter by about 1.82 times when the number of cores increased from two to four. On eight cores, it is about 1.7 times faster than on four cores. Similar scaling behavior was also observed with Δ*t* = 0.2 ms. On eight cores, pSpatiocyte (Δ*t* = 0.5 ms) is roughly 55 times faster than the parallel version of ReaDDy. It also required about 45- and 14-fold shorter runtimes than Smoldyn and Spatiocyte, respectively to complete the simulation on the workstation. Overall, the benchmark results show that pSpatiocyte has a significant performance advantage over other well-known microscopic particle simulators.

### Parallelized simulation of MAPK model

In the dual phosphorylation-dephosphorylation cycle of the MAPK cascade, shown in Figure 7A, molecular rebinding effects at the microscopic scale can alter the macroscopic dynamics of the system [36]. MAPK kinase (KK) phosphorylates MAPK (K) in a two-step process to generate a doubly phosphorylated MAPK (Kpp), whereas the phosphatase, P dephosphorylates Kpp twice to recover K. Upon unbinding from their products, the enzymes go through an inactive state (denoted by ^*^). The time required to reactivate the enzymes is given by *τ*_*rel*_ ≃ 1*/k*_*a*_, where

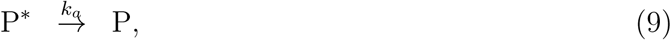

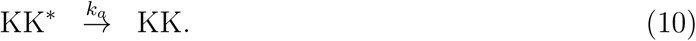

**Figure 7:**
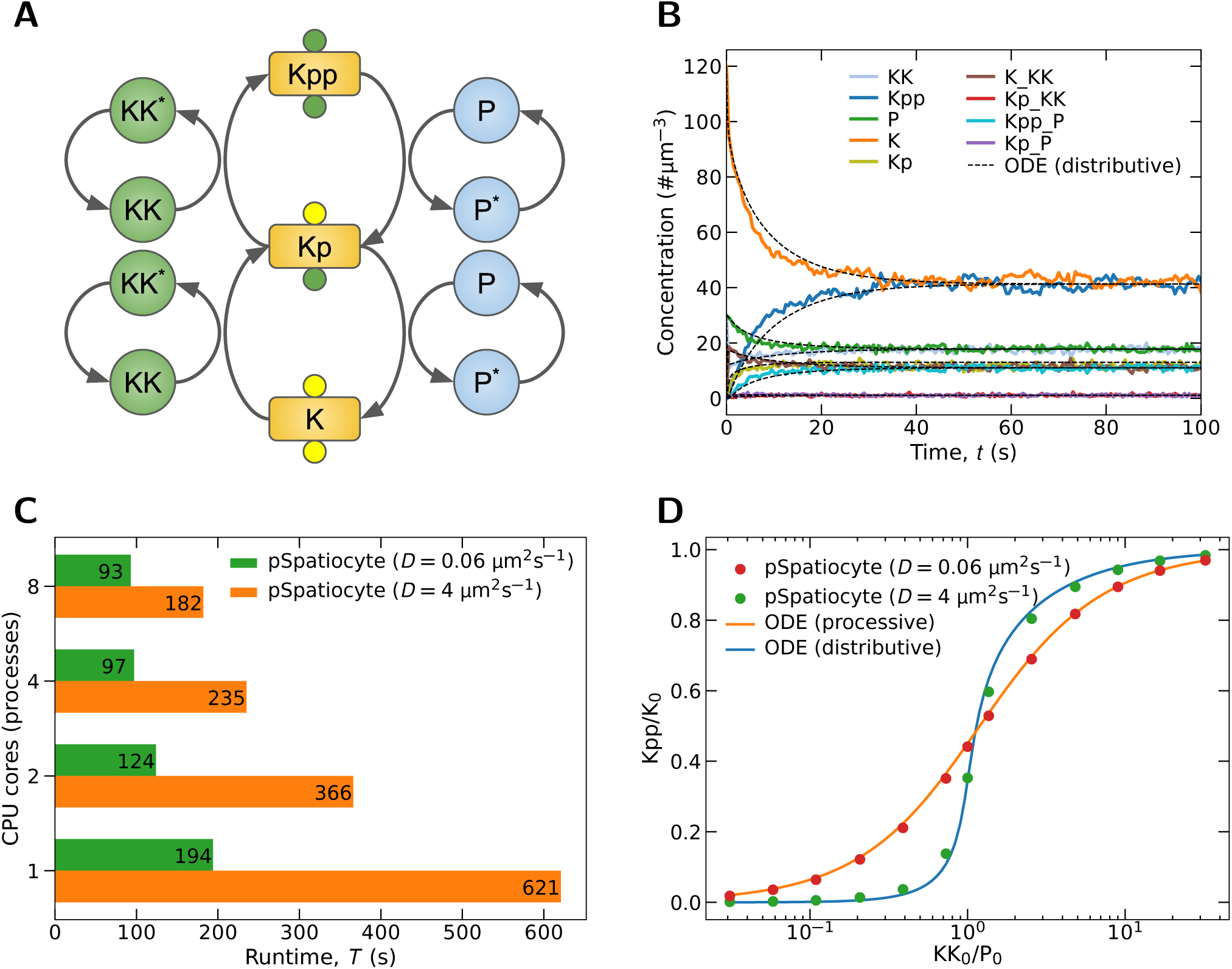
Parallelized simulation results of MAPK model. (**A**) Dual phosphorylation-dephosphorylation cycle of the MAPK cascade. (**B**) Concentration profiles when KK_0_*/*P_0_ = 1 and *D* = 4 *µ*m^2^s^−1^. Simulation was executed with all eight-cores of the workstation. (**C**) Scalability of pSpatiocyte at low molecule number. Shown are the runtimes of pSpatiocyte when simulating the MAPK model (KK_0_*/*P_0_ = 1) for 10 s with varying number of cores and diffusion coefficients. The model consists of 1800 total molecules. (**D**) Response curves for different diffusion coefficients. All simulations were executed with all eight cores of the workstation.

If the enzyme-substrate reactions are diffusion-limited and *τ*_*rel*_ is short, a newly dissociated enzyme can rebind to its product to catalyze it again, before escaping into the bulk. Takahashi et al. [36] have previously shown with eGFRD particle simulations that these rebinding events can change the response sensitivity of the phosphorylation state, which could result in the loss of bistability. We have also recently replicated the results with Spatiocyte [43]. These spatiotemporal correlations between enzyme and substrate molecules, and fluctuations at the molecular scale are difficult to be captured by RDME and PDE-based methods.

As an example of pSpatiocyte biological application and to further verify the method, we have simulated the MAPK model with the same parameters from [36] but increased the volume tenfold (10 *µ*m^3^) to raise the computational cost. The initial molecule numbers of K, KK and P were 1200, 300 and 300, respectively. There were no initial molecules for the remaining species. The model was simulated for 300 s with *τ*_*rel*_ = 1 *µ*s. For all species, *D* = 4 *µ*m^2^s^−1^. The lattice size was 476^3^ with *r*_*v*_ = 2.5 nm. Figure 7B shows the concentration profiles in the first 100 s of the simulation. The total runtime of the simulation was 5466 s using all eight cores of the workstation. In contrast, it took 76650 s to complete the simulation with Spatiocyte. Thus, pSpatiocyte is about 14 times faster than its sequential counterpart. As comparison, in Figure 7B we have also plotted the predictions of the corresponding mean-field (ODE) MAPK model [36]. The model describes the diffusion of molecules implicitly by renormalizing the reaction rates. pSpatiocyte concentration profiles coincide well with the ODE outcomes, although in the former, there were fluctuations from the small number of reacting molecules.

The total number of molecules in the MAPK model is 1800, which is a small number for high-performance simulations. To evaluate how well pSpatiocyte scales with such low number of molecules, we compared the runtimes with varying number of cores and diffusion coefficients. Figure 7C shows the runtimes when the model was executed for 10 seconds. With *D* = 0.06 *µ*m^2^s^−1^, pSpatiocyte is about twofold faster on four cores than on a single core. Increasing the number of cores from four to eight did not noticeably improve the runtime further. However, with *D* = 4 *µ*m^2^s^−1^, the speedup achieved with four cores is 2.6 times. With the addition of another four cores, the speedup increased to 3.4 times. This is notable because pSpatiocyte is still able to achieve a favorable speedup although each core only executed on average 225 molecules. Compared to the slower diffusion model, the shorter diffusion interval 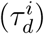 when *D* = 4 *µ*m^2^s^−1^ increases the computational cost more than the communication overhead, resulting in improved speedups.

We also simulated the MAPK model with different initial ratios of KK_0_*/*P_0_ and diffusion coefficients, and obtained the steady-state Kpp*/*K_0_ curves as shown in Figure 7D. The outcomes of the corresponding mean-field models of a distributive (with *D* = 4 *µ*m^2^s^−1^) and a processive (with *D* = 0.06 *µ*m^2^s^−1^) system are also shown. In the distributive scheme, the enzyme needs to detach from the substrate before it can catalyze it the second time. The double encounters between enzyme and substrate molecules can lead to ultrasensitive switchlike response. Conversely, in the processive scheme, a single encounter between them is sufficient to generate the dual modifications of the substrate. At fast diffusion (*D* = 4 *µ*m^2^s^−1^), pSpatiocyte response curve agrees well with that of the distributive mean-field model. However, at much smaller diffusion coefficient (*D* = 0.06 *µ*m^2^s^−1^), it instead reproduces the graded response curve of the processive model.

How slower diffusion in the pSpatiocyte model weakens the switchlike response curve can be explained as follows. At *D* = 4 *µ*m^2^s^−1^, the resulting diffusion interval, 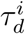 and walk event interval, 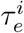 for the enzymes are the same (1 *µ*s) because the walk probability, *α*_*i*_ = 1. Since 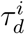 and *τ*_*rel*_ have the same intervals, after catalyzing the substrates the first time, the enzymes KK^*^ and P^*^ can escape into the bulk before they can reactivate and rebind with the substrates. In constrast, at *D* = 0.06 *µ*m^2^s^−1^, it gives 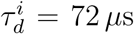 and 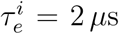 for the enzymes because *α*_*i*_ = 0.028. Since 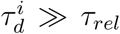, the enzymes have enough time to rebind with their substrates upon reactivation, before they can escape. This processivelike mechanism leads to the graded response curve as shown in Figure 7D. Overall, we note that pSpatiocyte correctly reproduces the expected ultrasensitivity dynamics of MAPK [36].

## CONCLUSION

We have developed a high spatiotemporal resolution parallel stochastic method to simulate intracellular reaction-diffusion systems on HCP lattice. To realize large-scale parallel computations, we have introduced several advanced simulation schemes, including a twisted Cartesian coordinate system with pointer-free voxel access, a parallelized event scheduler with priority queue, synchronized random subvolume executions and parallelized Gillespie’s direct-method using globally seeded random number generators, and three-stage inter-process data transfers.

We have also validated the physical correctness of the simulator. The simulated diffusion rates in dilute conditions showed very good agreement with theory. In crowded conditions, the diffusion rates decreased, as expected. Further work is required to compare the crowded diffusion behavior on lattice with the relation between diffusion and excluded volume fraction obtained in continuous space [68]. Both irreversible and reversible reaction curves coincided very well with predicted ODE results. Parallel performance of diffusion on the K computer was sufficiently high for large-scale computations. From the viewpoint of strong scaling, pSpatiocyte achieved a 7686-fold speedup with 663552 cores compared to the runtime with 64 cores on a 2048^3^ lattice. The efficiency was equivalent to 74.1%. In terms of weak scaling, efficiencies of at least 60% were obtained.

In the Michaelis-Menten enzymatic reaction benchmark, pSpatiocyte performed significantly better than other well-known microscopic particle simulators. On a workstation with eight CPU cores, pSpatiocyte is about 55 times faster than the parallel version of ReaDDy and 45 times faster than Smoldyn. In addition, the parallelized simulation of the MAPK model revealed that the program can correctly capture the weakening of ultrasensitive response by enzyme-substrate rebindings at very short timescales. The accurate simulation of the model also demonstrated that pSpatiocyte is applicable in real biological problems. On the same workstation with eight cores, pSpatiocyte required 14-fold faster execution times than the sequential version of Spatiocyte to simulate the MAPK model. Notably, pSpatiocyte is able to achieve 3.4 times speedup with all cores on the workstation although the average number of molecules executed per core is only 225.

In recent papers by Smith and Grima [72] and Novak et al. [73], RDME reactions with non-mass-action propensities such as Hill-type and Michaelis-Menten were shown not converging to the chemical master equation. At present, both sequential and parallel versions of Spatiocyte only support elementary reactions, namely unimolecular and bimolecular reactions. Since all complex reactions, including Hill-type and Michaelis-Menten, can be broken down to elementary (mass-action) reactions, the currently supported reactions should be sufficient for most modeling purposes. Nonetheless, for the convenience to directly model complex reactions, it would be beneficial to support them in the future. Further work would also be needed to solve the convergence problem.

Recently, several GPU-based high-performance simulators of reaction-diffusion systems have been reported [46, 48, 50–52, 74]. The ability to run fast simulations on common workstations equipped with GPUs supports wider application of the simulators. Implementing the Spatiocyte method to run on GPUs should be straightforward since most of the parallelization schemes presented in this work can also be applied on a GPU.

## ABBREVIATIONS

1D: one-dimensional
2D: two-dimensional
3D: three-dimensional
CPU: Central processing unit
FLOPS: Floating point operations per second
GPU: Graphics processing unit
HCP: Hexagonal close-packed
MAPK: Mitogen-activated protein kinase
MD: Molecular dynamics
MPI: Message passing interface
MSD: mean-squared displacement
PDE: Partial differential equations
RDME: Reaction-diffusion master equation
SCK: Smoluchowski-Collins-Kimball

## ACKNOWLEDGMENTS

We thank Steven Andrews and Frank Noé for their help with the benchmark models of Smoldyn and ReaDDy, respectively. We are grateful to the four anonymous reviewers for their constructive comments. We thank Peter Karagiannis and Kylius Wilkins for reading the initial version of the manuscript and providing valuable suggestions. AM wishes to thank Yukihiro Eguchi for continuous encouragement and Hisashi Nakamura of RIST for giving him the opportunity to start this study.

## AVAILABILITY OF DATA AND MATERIALS

The pSpatiocyte software and the complete scripts to generate the data in this study are available at https://github.com/satya-arjunan/pspatiocyte.

## AUTHORS’ CONTRIBUTIONS

AM, KT and SNVA conceived the project. AM and SNVA conceived the parallelization schemes and simulation algorithm. AM and KI wrote the initial version of the simulator. SNVA wrote, tested and optimized the simulation algorithm and software, and created the build system. AM and SNVA generated data and wrote the paper. All authors read and approved the final manuscript.

## ETHICS APPROVAL AND CONSENT TO PARTICIPATE

Not applicable.

## CONSENT FOR PUBLICATION

Not applicable.

